# Criticality in the conformational phase transition among self-similar groups in intrinsically disordered proteins: probed by salt-bridge dynamics

**DOI:** 10.1101/2020.03.30.016378

**Authors:** Abhirup Bandyopadhyay, Sankar Basu

## Abstract

Intrinsically disordered proteins (IDP) serve as one of the key components in the global proteome. In contrast to the dominant class of cytosolic globular proteins, they harbor an enormous amount of physical flexibility and structural plasticity enforcing them to be retained in conformational ensembles rather than well defined stable folds. Previous studies in an aligned direction have revealed the importance of transient dynamical phenomena like that of saltbridge formation in IDPs to support their physical flexibility and have further highlighted their functional relevance. For this characteristic flexibility, IDPs remain amenable and accessible to different ordered binding partners, supporting their potential multi-functionality. The current study further addresses this complex structure-functional interplay in IDPs using phase transition dynamics to conceptualize the underlying (avalanche type) mechanism of their being distributed across and hopping around degenerate structural states (conformational ensembles). For this purpose, extensive molecular dynamics simulations have been done and the data analyzed from a statistical physics perspective. Investigation of the plausible scope ‘selforganized criticality’ (SOC) to fit into the complex dynamics of IDPs was found to be assertive, relating the conformational degeneracy of these proteins to their multi-functionality. In accordance with the transient nature of ‘salt-bridge dynamics’, the study further uses it as a probe to explain the structural basis of the proposed criticality in the conformational phase transition among self-similar groups in IDPs. The analysis reveal scale-invariant self-similar fractal geometries in structural conformations of different IDPs. Also, as discussed in the conclusion, the study has the potential to benefit structural tinkering of bio-medically relevant IDPs in the design of biotherapeutics against them.

## 1. Introduction

Complex systems^1^ exist in *nature* where the structure of the system, in an abstract sense, degenerates among some population ensembles of various conformations [1]. Here structure may refer to the structural conformations in quantum particles [2,3], isomerism in stereo-chemistry, rotameric variation in amino acid side-chains [4] and to wherever the concept of degenerate states in structural ensembles may be applicable. Even the synchronization pattern of electrical activities of neurons in different parts of the brain [5] or pattern of economic consumption among different social groups [6] may be mapped to an abstract structural ensemble consisting of degenerate states. Structural degeneracy is important as it provides the system the flexibility to exhibit different properties, switch between different *modus operandii,* which, in the context of living systems (or functional unit of a living system, say, proteins), supports a variety of housekeeping as well as additional functionalities.

Proteins serve as the prime functional biomolecule of life *per se.* Their functions vary across a wide range from serving as enzymes in biocatalysis, signal transducers, transporters, molecular motors, providing elasticity to soft tissues like hair (fibrous proteins like keratin), tensile strength as well as flexibility to the muscle (collagen) [7], in the essential construction of the cytoskeleton (lamin [8]), acting as channels, gateways, molecular filters (membrane proteins [9]) and many more. Other than the special class of anchored (i.e., membrane proteins) and fibrous proteins, they generally remain in the cytosol as molecular globules (globular proteins) engaged in a certain type of fold (i.e., their 3D structural getup) consistent with their specific (routine) function. This gives birth to the classical view of protein folding consistent with the ‘sequencestructure–function’ paradigm. In contrast, there are multi-functional proteins as well, wherein, different parts of the protein 3D structure (e.g., domains, active and allosteric sites) serve to implement the few functions they are evolved to deliver. While, these molecular evolutionary strategies serve to construct the general rules of the protein structure-function paradigm, another variant has recently been discovered in the protein world, namely, fold-switch proteins [10] which switch between (a few) folds to support more than one functionality, generally induced by their chemical environment [11]. However, in all these cases, the notion of multi-functionality only varies within a few types of (premeditated) functions structurally not allowing the scope to get involved in sudden emergency *ad-hoc* functionalities as may be required contextually. Along with the increasing complexity in living systems with time (especially relevant for the modern human race) the demand of multi-functionality has increased in a proportionate manner. Intrinsically disordered proteins (IDP) [12], yet another relatively recent member of the protein family, a product of ever-increasing micro-evolutionary stress, has been revealed, unmistakably to have the potential to perform a various kind of functions to serve in such complex scenario [13], even unprecedented at times, characterized by their ‘unusual’ and ‘mysterious (meta) physics’ [14–17]. This great functional potentiality in these molecules is due to their inherent structural plasticity [18] and enormous amount of physical flexibility [18,19], characterized by their being present in conformational ensembles rather than one or two single fold(s) [20] throughout their entire life-span. Such conformational ensembles are further characterized by a population of structurally degenerate states, transitions between which are found to be ranging in tenths of *Å* to *nm* in length and *ns* to *s* in time-scale in proteins. Such multi-scale phenomena are essential for the proposed multi-functionality (even in the case of globular proteins, engaged in more than one function, say, allosteric signaling). Hence, it is important from a theoretical as well as a medical perspective to understand the phase transition phenomena among different degenerate states in intrinsically disordered proteins. In this paper, different degenerate states (represented by average temporal^2^ structures) of a collection of intrinsically disordered proteins are captured from molecular dynamics simulation (MDS) data using clustering analysis. This was followed by the implementation of statistical physics and phase transition dynamics to capture the equilibrium populations of the states, wherein, a major emphasis has been put to ‘criticality’ which potentially relates to conformational degeneracy and multi-functionality of these proteins.

The criticality hypothesis in phase transition dynamics refers to a system that may be poised in a critical state at a boundary between different dynamic manifolds characterized by its phase. Selforganized criticality (SOC) refers to the property of a large-scale dynamical systems where critical points also play the role of an attractant, such that the collective behavior shows some invariant characteristic of phase transitions in terms of the critical point without controlling parameter values. These types of systems are found to exhibit marginally stable behavior, effectively tuned towards criticality itself as it evolves, wherein, the event size obey a characteristic power-law distribution [21–24]. The example of SOC ranges from the simple geophysical phenomenon of piling sand to sophisticated phase transition in neural dynamics and brain functioning mechanism. The typical example of SOC is founded in non-equilibrium nonlinear systems with high degree of freedom. The concept was first introduced by Bak et. al., in a paper [21] in 1987 and is well accepted as a possible mechanism of emergence of complexity in nature. This was followed by some studies by Tang and Bak on scaling relation [22], mean field approximation [23] for SOC and relation of complexity with criticality [24]. Quickly these concepts were successfully applied across several fields of complex dynamics such as geophysics, Plasma physics and cosmology, quantum gravity, sociology, ecology, evolutionary biology, neurobiology [25–27], economics, optimization, bio-inspired computing [28] and many others. Consequently the concept of SOC was applied to several other fields of natural complexity, which has been already evident for emergence of scale-invariant behaviors in large-scale physical or social systems. SOC was found to be successful to explain and analyze several complex systems and phenomenon like earthquakes, landscape formation, forest fires, solar flares, landslides, epidemics, fluctuations in economic systems such as financial markets, neuronal avalanches in cortex [26,29], biological evolution, I/f noise in the amplitude envelope of electrophysiological signals [25] etc. These studies on SOC include attempts to model the dynamics as well as extensive data analysis to determine the characteristics and condition for existence of natural scaling laws. Also several recent studies have shown to evolve scale-free networks as an emergent phenomenon in SOC [30]. On the contrary, other researches on the solvent-accessible surface areas of globular proteins suggest that SOC exist independently of any physical space or dynamics [31]. Also quantification of the differential geometry of proteins from the SOC in its structural dynamics resolves many unsolved questions regarding the biological emergence of complexity [32].

In this paper, we focus on the characterization of criticality in the context of conformational phase transition in IDPs based on their time-evolved atomic coordinates (i.e., Molecular Dynamic simulation trajectories). This analysis is further coupled and complemented by the study of salt-bridge dynamics – which has already been revealed [33] to serve as a meticulous mechanism to contribute to (and retain) the characteristic physical flexibility in these proteins, abundant in charged amino acids. Taking an important bold step forward, the current study attempts to use ‘salt-bridge dynamics’ as a probe to investigate (and reveal) the scope of the aforementioned criticality in the conformational phase transition among self-similar groups in IDPs. To that end, the study should potentially serve as an essential footstep in the plausible control of protein functionality and in the design of biotherapeutics particularly relevant for neuro-degenerative disorders given rise by malfunctioning IDPs [34,35].

## 2. Materials and Methods

### 2.1. Selection of IDPs

Idp’s chosen for the current study were kept precisely the same as that of an earlier study [33], wherein, four proteins were chosen with their degrees of structural disorder varying from 43 to 100% in their native states. Two partially disordered proteins (IDPRs), namely, the scaffolding protein GPB from Escherichia virus phix174, (PDB ID: 1CD3, chain ID: B) and the human coagulation factor Xa, (PDB ID: 1F0R, chain ID: B) along with two completely disordered proteins (IDPs), namely, α-synuclein (α-syn) and amyloid beta (Aβ42) were chosen to construct the set. The sequences of the IDPs were obtained from the DISPROT database [36]. For 1CD3, 1F0R their X-ray structures (resolution: 3.5 Å & 2.1 Å respectively) were obtained from the Protein Data Bank (PDB) [37]. The missing coordinates corresponding to the disordered regions were identified by comparing the SEQRES and ATOM records in their corresponding PDB files. The final atomic models were obtained from the earlier study after the disordered regions in 1CD3, 1F0R along with the full-length sequences of the completely disordered proteins (α-syn, Aβ42) were built using MODELLER [38].

### 2.2. Molecular dynamic simulation

As an improvement to the earlier studies [33,39], an altogether different new protocol was adapted for performing the explicit-water Molecular Dynamics (MD) simulation for the chosen IDPs, wherein the production phase was made to run for a 5-fold longer period of time (500 ns in contrast to 100 ns in the earlier studies) in GROMACS v.2018.1 [40] using the latest available force-field GROMOS96 54a7 [41] associated with the MD package (in contrast to ff99SB [42,43] in AMBER 12 [44] opted for the earlier studies). Periodic boundary conditions were used with the SPC [45] water model recommended for all available GROMOS 96 force-fields. Solvation and charge neutralization of the proteins were subsequently followed by two rounds of energy minimization (in staid of just one round used in earlier studies [33,39]) using the in-built minimizer module within GROMOS 96. In the first round, 200 steps of the relatively much faster steepest descent method was used wherein atoms are moved so as to reduce the net forces on them leading to an instantaneous freezing of the system. This was followed by 19800 steps of the more productive conjugate gradient method to remove unfavorable steric interactions. The energy minimized protein – solvent system was then equilibrated in an NVT ensemble followed by an NPT ensemble for 100 ps and 5 ns respectively. The initial temperature set for the NVT ensemble was 100 K which was gradually raised to 300 K at constant volume and was kept the same for the entire NPT equilibration while the pressure maintained at 1 bar. The production run of the MD simulation was done in an NPT mode for 500 ns with a time step of 2 fs for each equilibrated protein – solvent system. To maintain constant temperature, Berendsen’s temperature bath was used with a coupling constant of 2 ps, while barostat with a coupling constant of 1 ps was used to regulate the constant pressure. Trajectories were written at an interval of 2 ps, resulting in 2,50,000 frames (or time-stamps). All analyses were performed on the post-equilibrium 500 ns long trajectories (for all four proteins).

### 2.3. Identifying Salt-bridges

As is standard in protein-science literature [33,46,47], ionic bonds within IDPs were detected when a positively charged nitrogen atom from the side-chains of lysine, arginine or positively charged histidine were found to be within 4.0 Å of a negatively charged side-chain oxygen atom of glutamate or aspartate.

### 2.4. Clustering of MD Simulation data based on RMS distances among the snapshots

Let the simulated MD trajectories be split from time-step 1 to time-step *n + n_t_*, wherein, the first *n_t_* steps are considered to be in transient phase and are removed to get the structural conformations in the final n steps. Let, A be an *n×n* matrix where *A* = (*a_ij_*)_*n×n*_ where *a_ij_* are the root mean square (RMS) distances between the conformations obtained at *i^th^* and *j^th^* time-steps. Also, let, B be the associated adjacency matrix where the structures at *i^th^* and *j^th^* time-steps are connected (or adjascent), if their distance is found to be less than a pre-defined threshold (*θ*). Furthermore, the structures considered as nodes of the adjacency matrix were sorted according to their degree (i.e., the number of structures connected to a particular structure or node). Let *I* be the sorted list of structures, *CC* be the list of representative (average) structures defined as the cluster centers and *CN* be the list of cluster numbers for each structure denoting the cluster in which the structure belongs to. In the present study, θ was set to 5Å and 7.5Å respectively for 1CD3 and the rest of the three IDPs, considering their relative abundance (and scarcity) of ordered secondary structural (particularly helical) content (see section **2.1. Selection of IDPs**) coupled with previous knowledge of relative instability of the proteins [33,39].

The adjacency matrix B can be represented mathematically as the following.

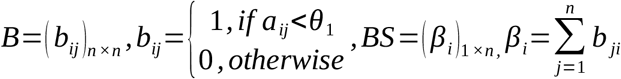

The clustering is done by the following algorithm.

*Push first element of I into CC*

*For step i=2 to n*

> *Push I(i) into CC if distance between all structures already in CC and the structure at time step I(i) be greater than θ.*

*Generate m cluster centers.*

*For step i=1 to n*

> *Each structure is assigned a cluster number according to the cluster center that lies at the minimum distance from the structure,*
>
>
> 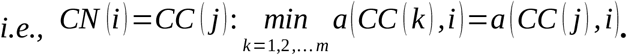

Now, the transition matrix *TM* is generated such that *TM* = (*tm_ij_*)_*m×m*_ where *tm_ij_* represents the number of times the structure was found in *CC*(*i*) at the previous time-step and in *CC*(*j*) at the next time-step.

The transition probability matrix *M* is obtained by dividing every element of *TM* by its corresponding row sum, i.e. 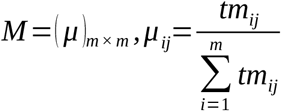. These *m* structures in *CC* are considered to be *m* different average structures or states of the time-evolved biomolecule.

### 2.5. Efficiency of clustering

The aforementioned adjacency matrix *B* can be interpreted in terms of a graph, *G*=(*V,E*) having *V as its* set of vertices and *E as its* set of edges. The clustering efficiency of *G* could be determined by the clustering coefficient computed from *B* in the following way.

In the graph *G,* if the vertices *v_i_* and *v_j_* are connected by an edge, then the corresponding element of *B, b_ij_* will be 1 or 0 otherwise. The neighborhood *N_i_* of vertex *v_i_* is defined as its immediately connected neighbors as *N_i_*=|*v_j_*:*b_ij_*=1, *i,j*=1,2,…,*n*]. Also, let, *k_i_* denote the number of vertices in the *i^th^* neighborhood, i.e. |*N_i_*| = *k_i_*. The local clustering coefficient *C_i_* for a vertex *v_i_* is then given by the proportion of links between the vertices within its neighborhood divided by the total number of links that could possibly exist between them. For an undirected graph *G*, the local clustering coefficient of its i^th^ node, *C_i_* can then be defined as follows.

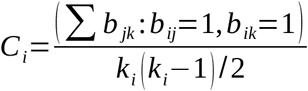

All *C_i_*’s in a graph can further be averaged to return the average clustering coefficient (*C*) of the graph.

Also, the edge density within the clusters and that in between different clusters gives a good estimation of clustering efficiency of any clustering scheme. We define an ordered parameter *op* as the ratio of the intra-cluster edges and the total number of edges, i.e.

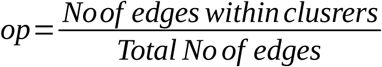

*op* will trend towards 1 as the clustering scheme becomes better in the sense that a greater proportion of edges become intra-cluster.

### 2.6. Phase transition

In the context of the time-evolved structures of IDPs (obtained from their molecular dynamic simulation trajectories), several self-similar conformations could be considered as different structural phases of these molecules and their intrinsic disorder could be explained as the transition dynamics among these different phases. In this paper, self-similarities in structural conformation of IDPs are grouped on the basis of their relative deviations and is aided in the formation of the conformational clusters which could be interpreted as structural phases of these biomolecules. Hence, the intrinsic disorder of IDPs could be explained by the phase transition dynamics among *m* different phases which could be parameterized by the transition probability matrix M directly obtained from the MD simulation trajectories.

Let, *p*(*i,t*),*i* = 1,2,…,*m* be the probability that the biomolecule is in structure *CC*(*i*) at time *t* and *P*(*t*)=(*p* (1,*t*), *p* (2, *t*),…. *p* (*m,t*))^*T*^. Then the phase transition dynamics could be written as follows:

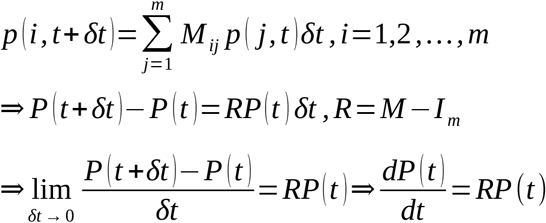

*I_m_* is the identity matrix of order *m*. Row sum of each row of *M* is *1*, thus *R* is a matrix with zero row sums. Hence *R* has a fixed Eigenvalue zero and the stability of the dynamics is determined by its largest Eigenvalue. Also,

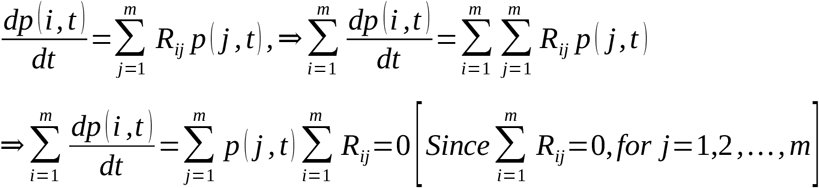

Hence, the sum of all probabilities remain unchanged and *P* forms a simplex in phase transition between *m* states.

### 2.6. Analysis of salt-bridge persistence

Again, a similar protocol was adapted from the earlier study [33] with only minor contextual variations. As was done before, simulated conformations were collected at an interval of 50 ps across the entire 500 ns MD trajectory for each idp, resulting in 10000 protein conformations spanning the entire length of trajectory. Each trajectory was then split into clusters based on RMS deviation (as detailed above) and in each of these resulting clusters, the dynamic persistence (*pers*) of a particular salt-bridge was calculated as the ratio of the number of protein conformations to where the salt-bridge was found to form with respect to the total number of conformations in that cluster. Even a single occurrence of a salt-bridge in a given cluster was considered accountable in this analysis. Normalized frequency distributions of salt-bridge persistence were plotted for each cluster from these raw distributions (to be discussed in the next section). To figure out the representative ‘persistent’ salt-bridges in each cluster, a cut-off of 25% (i.e., *pers* ≥ 0.25: a salt-bridge found in at least 1/4^th^ of all sampled frames in a cluster) was considered optimum (as standardized in an earlier study [33]) and applied.

### 2.7 Test of Scale-freeness – signature of criticality

Scale-freeness indicates criticality [21,22] in phase transition, wherein, a system smoothly traverse around multiple unstable steady states. A system here could either be a physical or a chemical or a biological or any other complex system. For example, in the context of neural discharge of neurotransmeters, burst size is known to give signatures of scale-freeness in brain disorders (epilepsy for example) [27,29] as its distribution follows power-law with an appropriate exponent. To test an equivalent scale-freeness, an analogous analysis was adapted in the current study by studying the distribution of salt-bridge persistence in each cluster. No persistence cut-offs were used in this analysis, the entire numerical range of persistence (*pers*) values [0, 1] was binned in bin-size of 0.1 and normalized frequencies (*nf)* of salt-bridges with a given width of persistence (corresponding to a certain bin) were computed and plotted in a loglog plot (i.e., logarithm of normalized frequency in Y-axis as a function of persistence in X-axis: log(*nf*) vs. log(*pers*)). The bins with zero occupancy when converted to the log count gives a negative infinity (-Inf). For such instances, the ‘-Inf’ were replaced by contextually determined arbitrarily large negative finite values based on the range of obtained finite log(nf) values, for that given plot. Linear least-square fitting was performed on these experimental points, the R^2^ (coefficient of determination), fitting errors (root mean square deviation of the same) and the slope of the fitted straight-lines were recorded.

## 3. Result and discussion

### 3.1. Clustering: identification of conformational phases and phase transitions

The prime focus of the study was on the analysis of structural degeneracy of disordered proteins (from their MD simulation trajectories) and therein the characterization of conformational phases and transitions between them. Four representative IDPs, namely, 1CD3, 1F0R, α-syn and Aβ42 were selected for this purpose, the first two being partially and the next two being completely disordered. In the perspective of structural degeneracy of IDPs, it is important to identify the self-similarities of different structural conformations and club them into appropriate groups to classify all these conformations into few identifiable clusters. Here, different structural conformations of IDPs (as obtained from MD simulation) are categorized in several groups based on their dissimilarities that can be measured from their RMS deviations. These structural groups or clusters serve the purpose of characterization of the overall molecular dynamics in terms of few representative structural ensembles and are considered as dynamical phases in the molecular dynamics of corresponding IDPs. The phase transition dynamics are, hence, studied to analyze the persistence and long term behavior of the corresponding IDPs to retain them as a collection of representative structural conformations. The structural phases of these IDPs are thus derived as conformational clusters based on RMS distances between conformations by clustering analysis as elaborated in the Materials and Methods. The transition probability matrix for each IDP is extracted from MD simulation data. The phase transition dynamics are simulated for individual IDPs and the persistence of these phases or conformational clusters are analyzed therein. The fixed point or equilibrium in phase transition dynamics is obtained in terms of probability of attaining the aforementioned representative structures which could be interpreted as steady state persistence of each conformational cluster.

For 1CD3, three different structural conformations were found with their cluster centers obtained at time-stamp 154130, 249130 and 207505 where the corresponding equilibrium (or fixed point)in terms of probabilities (P) of attaining a representative structure were found to be 0.5433, 0.2585, and 0.1982 respectively. These probabilities can be presented as an array P (e.g., P=[0.5433, 0.2585, 0.1982]) with their elements adding up to 1. Similarly, for 1F0R, six different structural conformations were found with cluster centers at time-stamp 231130, 143755, 175380, 105130, 92505 and 214005 with the corresponding probability array as P=[0.3239, 0.1807, 0.1379, 0.1159, 0.1998, 0.0418]. For α-syn, five different structural conformations were found with cluster centers obtained at time-stamp 221505, 116255, 255005, 191630 and 135880 with P=[0.3706, 0.2145, 0.1556, 0.0857, 0.1736] while for Aβ42, six different structural conformations were obtained with cluster centers at time-stamp 210505, 120755, 190380, 134630, 216880 and 126005 with P=[0.3731, 0.1042, 0.0731, 0.2684, 0.1152, 0.0661].

Following the mathematical part of the clustering analysis, visualizations were subsequently done **(Figure 1, 2)** by (i) superposing the cluster centers (left panels of **Figure 1, 2**) leading to a reduced representation of the degenerate structural ensembles of the IDPs and (ii) also looking at them individually (right panels). The efficiency of clustering (or clustering efficiency) could be evaluated as the value of the ordered parameter (op) (see **Materials and Methods**) which has been derived and obtained as 0.5488, 0.8349, 0.7424 and 0.9506 respectively for 1CD3, 1F0R, α-syn and Aβ42. The average clustering coefficient (C) of the associated network (see **Materials and Methods**) gives a measure of how densely the clusters are packed. C was obtained to be 0.7836, 0.5818, 0.5948 and 0.5358 respectively for 1CD3, 1F0R, α-syn and Aβ42. From these numbers, it is evident that 1CD3 (among the four IDPs) has the least amount of structural degeneracy for having the lowest clustering efficiency (op) and the highest average clustering coefficient (C) in the whole set. That is to say that the structural conformations for 1CD3 are relatively more self-similar to each other. This is perhaps meaningful as 1CD3 is the protein that has the highest secondary structural as well as helical content (**Figure 1**) among the four IDPs (see **Materials and Methods**) and can therefore be interpreted as the closest (out of the four) to the class of globular proteins. Interestingly, the other partially disordered protein, 1F0R has far more structural diversity wherein the conformations are substantially different from each other, as reflected from its much lower clustering coefficient matching to the order to those obtained for the completely disordered proteins. Six different conformational phases describe the structural diversity in 1F0R and the relatively higher value of clustering efficiency (op) suggests that there is little structural resemblance among its conformational clusters or in other words the conformations within each cluster are more self-similar. These behavioral difference of the two partially disordered proteins, 1CD3 and 1F0R can also be interpreted from the perspective of the different type of salt-bridge formation in them, to be discussed in the next section.

**Figure 1.**
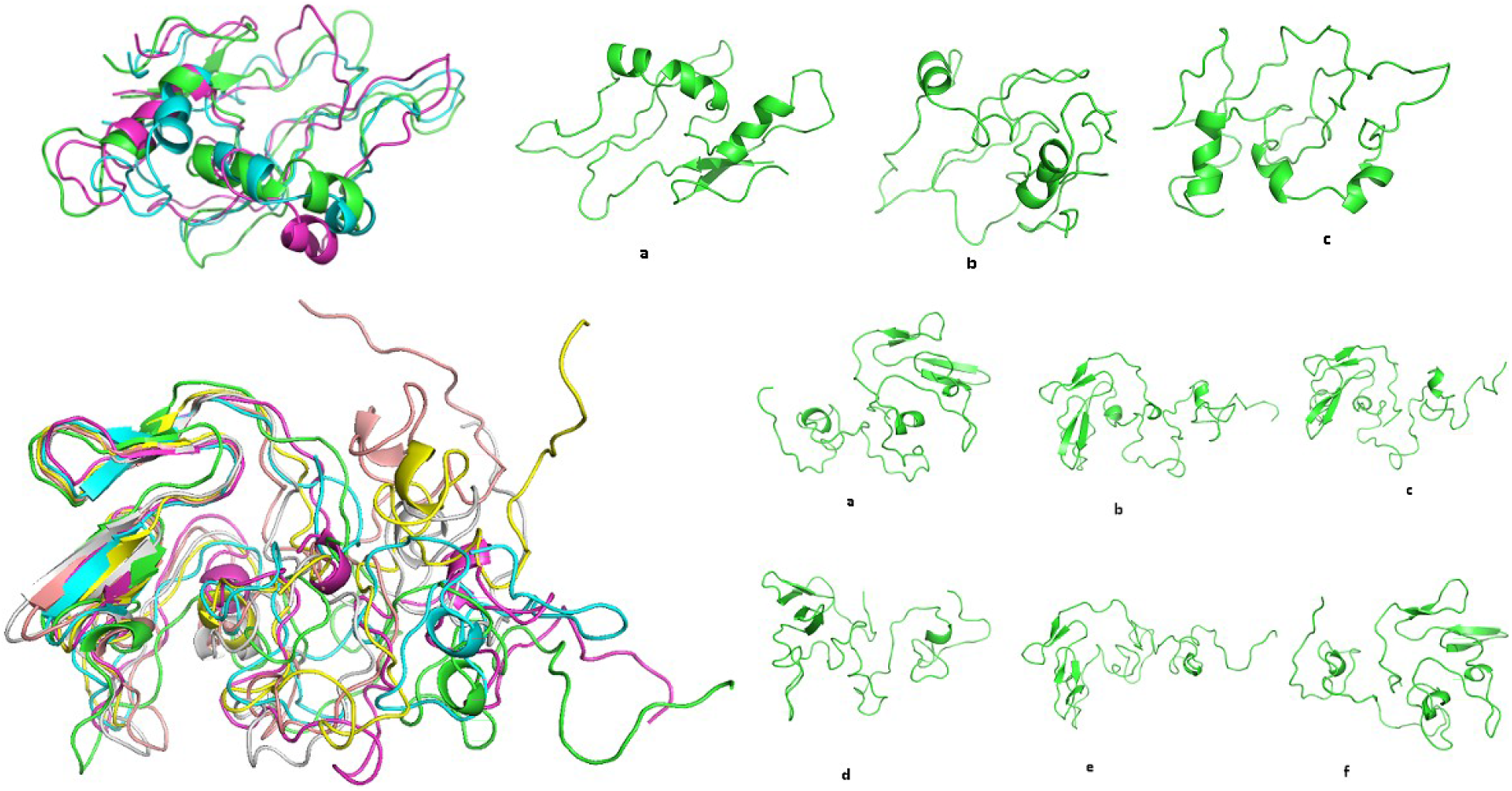
Upper Row: Structural conformations of 1CD3. The conformational clusters (phases) represented by the corresponding cluster centers: a) C1, b) C2, & c) C3 with time stamp 154130, 249130 & 207505 respectively presented individually on the right and superposed on the left. Lower Row: The same for 1F0R: a) C1, b) C2, c) C3, d) C4, e) C5 & f) C6 with time stamp 231130, 143755, 175380, 105130, 92505 & 214005 respectively.

**Figure 2.**
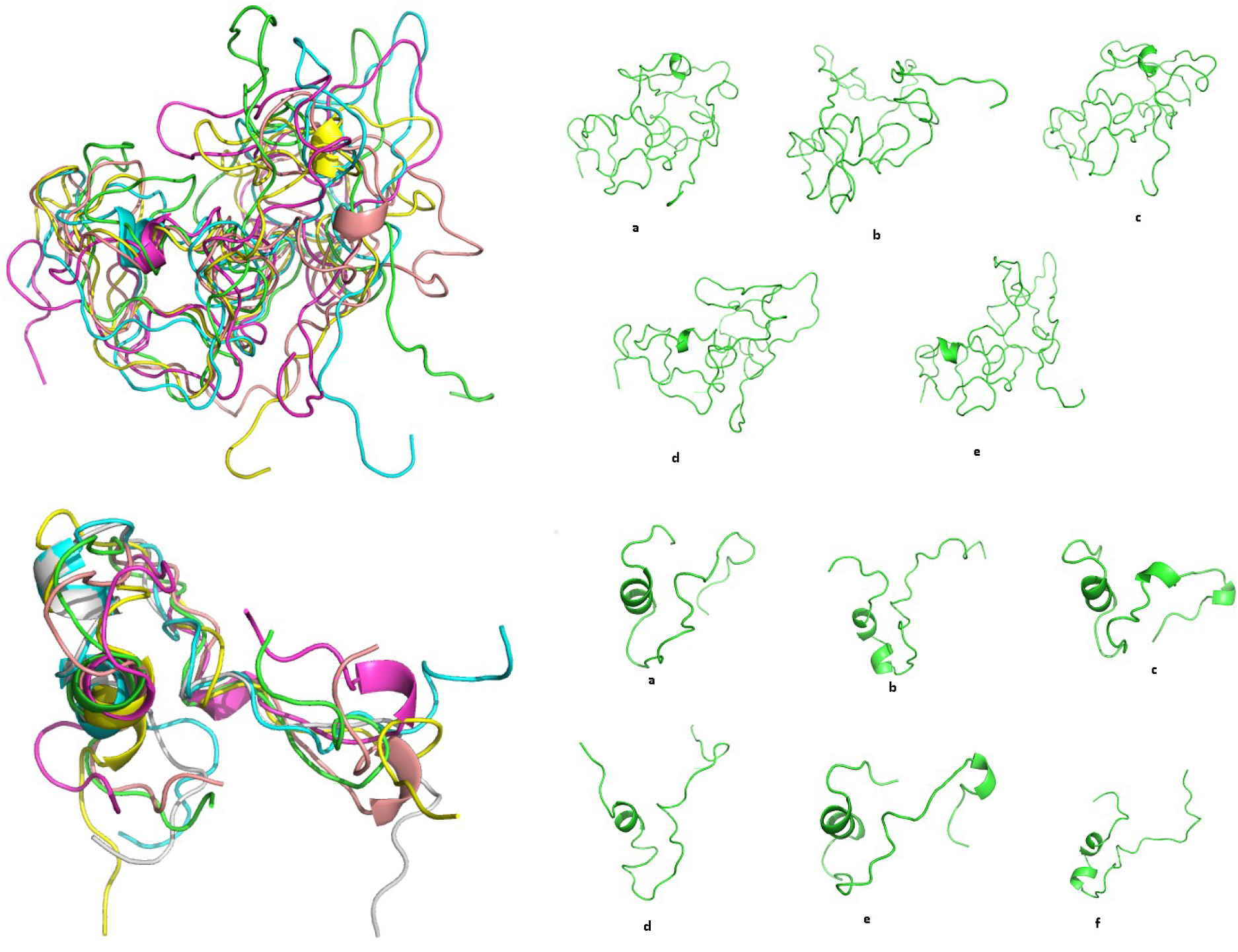
Upper Row: Structural conformations of α-syn. The conformational clusters (phases) represented by the corresponding cluster centers: a) C1, b) C2, c) C3, d) C4 & e) C5 with time stamp 221505, 116255, 255005, 191630 & 135880 respectively presented individually on the right and superposed on the left. Lower Row: The same for Aβ42: a) C1, b) C2, c) C3, d) C4, e) C5 & f) C6 with time stamp 221505, 116255, 255005, 191630 & 135880 respectively.

On the other hand, the completely disordered proteins, α-syn and Aβ42 naturally have substantial diversity in their structural conformations as reflected from their corresponding lower values of clustering coefficients. Among these two completely disordered proteins, Aβ42 has been classified into six conformational clusters (phases) which are fairly diverse from each other,wherein, the phases show a high degree of self-similarity within themselves (and higher than that of the other three proteins) as suggested by the highest value of its clustering efficiency (op) in the set. In case of α-syn, the value of clustering efficiency (op) is moderately high which suggests that α-syn also has fairly diverse structural conformations, wherein, the segregation of these conformations into five different phases are fare. These observations can be extended to infer that the molecular dynamics of α-syn is more continuous in nature than Aβ42 and the difference between self-similar groups in the former is relatively less. Overall, from this analysis we can conclude that 1CD3 has the most regular structure among the four, α-syn has a fairly diverse structure yet a continuous transition behavior, while 1F0R and Aβ42 both have proper diverse structural phases with substantial self-similarity among the conformational phases.

To understand the relative intensities of the conformational phases along time, the time-evolution of the probabilities of attaining different states or conformational clusters were plotted together (**Figure 3**). The equilibrium values of these probabilities were presented in the aforementioned array P. Here, it is interesting to find that ~55% of the population ensemble for 1CD3 solely represent its first conformation, C1. For 1F0R, the first two conformations, C1 and C2 add up to more than half of its population ensemble. Similarly, almost sixty percent of the population ensemble for α-syn map to two of its most populated conformations: C1 and C2 while more than sixty percent of the population ensemble for Aβ42 are distributed between C1 and C4. This give a nice structural insight into the conformational degeneracy of these IDPs which could be interpreted in terms of few representative conformations.

**Figure 3.**
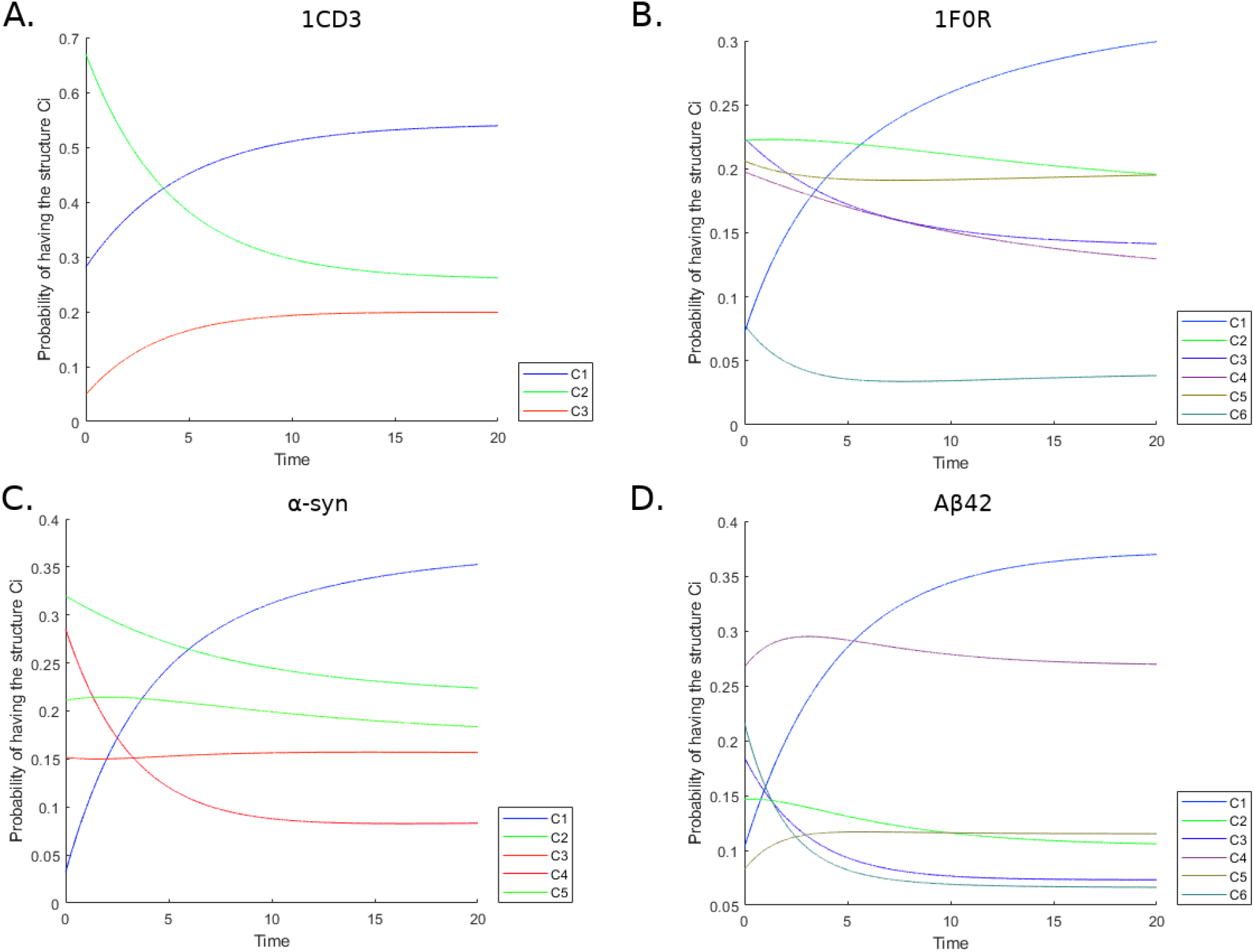
Simulated phase transition dynamics presented by the probabilities of attaining different states with respect to time for the four IDPs: A) 1CD3, B) 1F0R, C) α-syn & D) Aβ42. The simulations were done till equilibriation of the dynamics with a non-dimensionalized timescale. The time axis is presented in arbitrary units.

### 3.2. Transient Salt-bridge Dynamics

The transient nature of salt-bridge dynamics, or, in other words, the flitting character of ephemeral salt-bridges across the whole protein chain was previously revealed [33] to be crucial in retaining the conformational flexibility of disordered proteins. This, in turn, was found to be essential (at least *in-silico*) for such protein functions (in the binding of their globular partner proteins). From a mechanistic point of view, it was further revealed that these salt-bridges consisted of charged atom pairs continuously changing their ionic-bond partners and thereby collectively supporting different conformations. The mechanism thus functions as a ‘conformatonal switch’ in the context of idp-dynamics. Again, being evolved primarily as a ‘structural’ switch, the phenomenon has the potential to turn on another ‘functional’ switch from the ‘intra-’ to ‘inter-chain’ salt-bridges in the idp when its (globular) binding partner become available and accessible for binding. As a matter of fact, it is largely the short-lived salt-bridges of the idp (i.e., the ‘intra-chain’ salt-bridges) which collapses momentarily before concomitantly reuniting with charged groups coming from its globular partner (giving rise to formation of the ‘inter-chain’ ones) as they remain functionally far more open-ended and amenable as compared to the persistent salt-bridges of the idp. Overall, the transient salt-bridge dynamics in IDPs potentially serves as an initiation and stabilization mechanism for protein-protein binding in the context of an idp and its globular partner.

For the current study, salt-bridges were first identified in each conformational cluster and their persistence computed in that cluster. Likewise that of the earlier study [33] even a single occurrence of a salt-bridge in a cluster was considered important and recorded. The overall trends of the distributions of salt-bridge persistence (i.e., frequency vs. persistence) were found (**Figure 4**) reasonably similar to the one obtained for the entire trajectory [33], wherein, persistence bins above the cut-off of 0.25 (which was the previously standardized threshold to define ‘persistent salt-bridges’ [33]) were found to be roughly equally populated, followed by a long raised tail (left-peak in the plot) comprising of a high fraction of short-lived salt-bridges. The distributions visually resembled power series decays along the direction of increasing persistence and could best be fitted to rectangular hyperbola’s (y=k/x) where the proportionality constants (k) were determined based on the scale of the Y-axis. From the distributions, it was clear that, in each cluster, there were some persistent salt-bridges (potentially representative of that conformation) along with a large fraction of flitting salt-bridges over the whole chain, analogous to the invariant and variable parts of an equation respectively. It was also realized that to switch to another conformation (or conformational cluster) the protein has to undergo modulation in the two types of salt-bridges (persistent and ephemeral) at different degrees. In other words, for persistent salt-bridges, some of them may remain common or conserved between two or more conformational clusters, while the others may vary, and, the ubiquitous presence of the ephemeral salt-bridges across the dynamic protein chain may simply provide the matrix (acting as if like a buffer) to switch between conformations. That is to say that during this confromational switch, one or more persistent salt-bridges may break open and be replaced by other newly formed persistent salt-bridges while the ephemeral salt-bridges may simply rearrange themselves to fit the new conformation.

**Figure 4.**
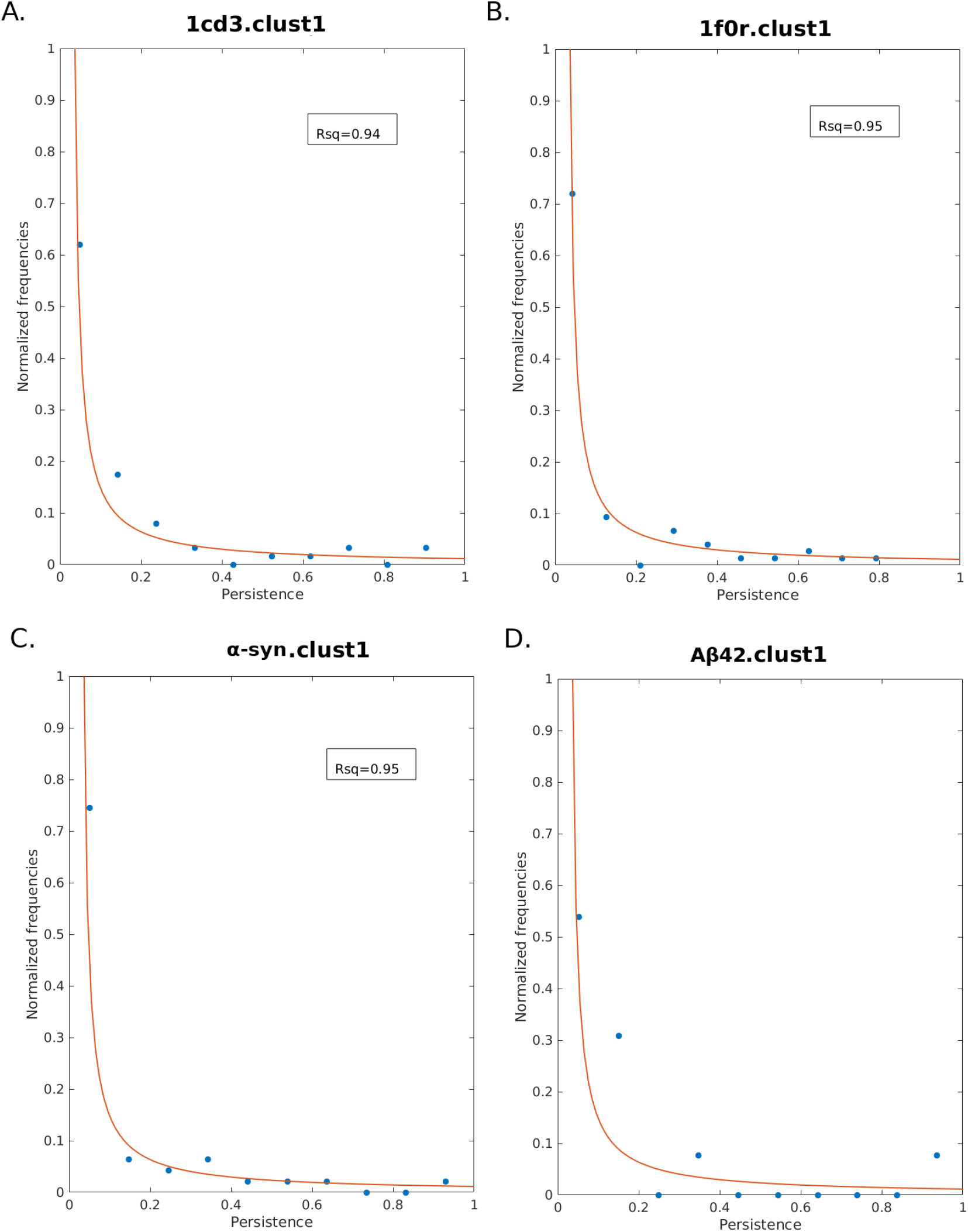
Distribution of salt-bridge persistence plotted for the first clusters (C1) of A) 1CD3, B) 1F0R, C) α-syn & D) Aβ42. The distributions could best be fitted to rectangular hyperbola’s.

The presence of the conserved salt-bridges across conformations (some of them even across the whole molecular dynamic trajectory) is meaningful and important, since, even IDPs in water (or in the cytosol) does not remain as completely extended elongated random coils or ‘ideal chains’^3^ rather undergo sequence dependent dynamic bending (primarily due to extensive electrostatic interactions throughout the whole chain). Hence, likewise the globular proteins, they too are physically restricted by some amount of local rigidity as imparted by short-range persistent saltbridges by often creating fairly stable loops and turns (sometimes even a short helix as is the case of Aβ42). Such constraints make them structurally approach porous globule – as was reflected from their shape factor profiles [33]. Broadly speaking, the short-range (sequence separation of less than ten amino acids) persistent salt-bridges thus could be envisaged essential for their basic time-evolved structural getup. On the other hand, the long or medium ranged salt-bridges across time can potentially give an idp its desired variation in structural identity across conformations, by preferring to form and remain persistent within certain conformational clusters while remaining absent in the others. The third prototype, a large abundance of the ephemeral saltbridges form and collapse momentarily in each conformational cluster, providing the necessary conformational entropy and flexibility required (even) within a cluster.

The Persistence vs. Contact Order profile (individually as well as when merged for all clusters, **Figure 5**) also largely followed a power series decay and could best be fitted to a rectangular hyperbola – meaning that the (high) persistent bins of salt-bridges were more populated with short-range than long-range contacts while the ephemeral salt-bridges had no such contact order preferences. To make a comprehensive test of the above hypothesis, the Contact Order (CO) vs. Persistence of salt-bridges were plotted individually for each protein twice: (i) for their full molecular dynamic trajectories (represented by black open circles in **Figure 5**) and (ii) for individual clusters (blue dots in **Figure 5**). Here, in this figure, it is to be carefully noted that points in **Figure 5** having the same abscissa (CO) and only differing within a narrow range of their ordinate (*pers*) actually correspond to the same salt-bridge. Among such a cluster of points, the encircled point correspond to the whole MD trajectory which only get split into different conformational clusters. We can consider the whole range of salt-bridges categorized into three classes (i) long range persistent salt-bridges (ii) short and moderate range persistent salt-bridges and (iii) ephemeral (i.e., short-lived) salt-bridges. The former class was found to be only little populated (occurred just twice for the two partially disordered proteins: 1CD3, 1F0R) while the later was heavily populated (with virtually no correlation with contact order) adding to the conformational entropy (as discussed earlier in this section). The second class having the highest X-width (i.e., persistence range) was of prime importance, wherein, persistence generally followed an inverse trend with respect to contact order. From a structural perspective, this class of salt-bridges appears to be potentially important for (a) creating small to moderate lengths of ‘loops and turns’ in the protein at different temporal phase and (b) remaining intact in/across conformation(s) (some of them even throughout the whole MD trajectory). They therefore impart a varying degree of local temporal structural constraints to the protein, which, in turn, collectively contributes to its unique 3D conformational getup corresponding to the cluster(s). It naturally follows that these may be envisaged as representative salt-bridges for the cluster(s).

**Figure 5.**
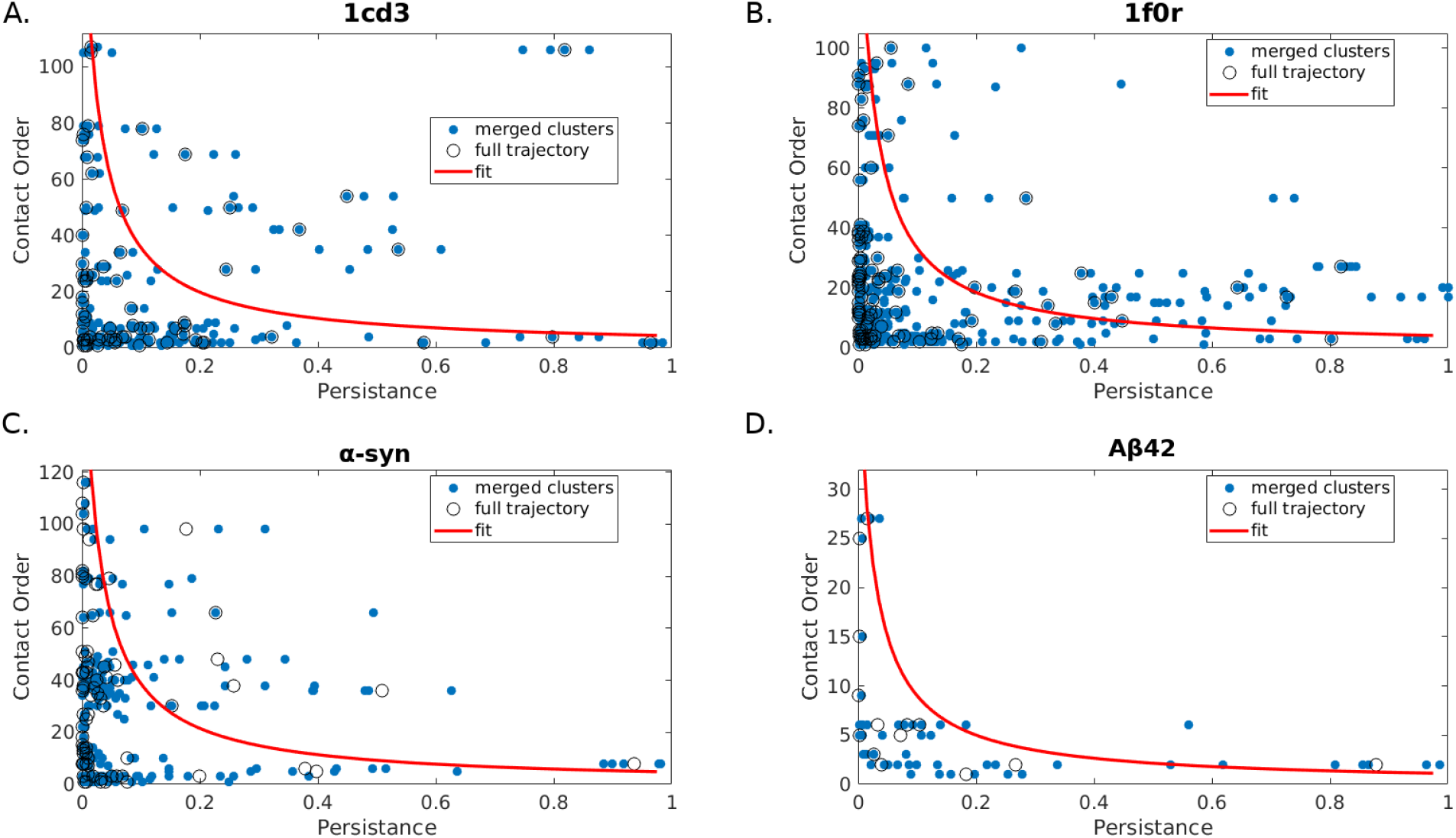
Persistence *(pers)* vs. Contact Order (CO) profiles for the IDPs. Persistence values for the full molecular dynamic trajectories are represented by black open circles while the same for individual clusters (i.e., ‘cluster persistence’) are presented as blue dots.

Looking closely into the two instances of long range persistent salt-bridges, they were found to represent two opposite end of the spectrum of possible molecular dynamic events. The one in 1CD3 (2-Glu ~ 108-Arg) had an overall persistence of 0.819 (the only encircled point in **Figure 5** panel A, right-top part of the plot) for the whole MD trajectory varying only from 0.747 to 0.864 among the three conformational clusters in the protein. Here we must recall the fact that the protein is partially disordered, having the highest percentage of secondary structural content: 56.7% [33]. From a detailed structural view (**Figure 6**), the salt-bridge was found to form between two anti-parallel beta strands coming from the two termini (N’ and C’-) which remain intact throughout the entire course of its dynamics, bringing and retaining the two end of the protein in close proximity and giving the protein its desired dynamic bending. The case therefore represents a salt-bridge mediated long range secondary structural association which is a conserved structural feature of the protein along the dynamics of its conformational variations.

**Figure 6.**
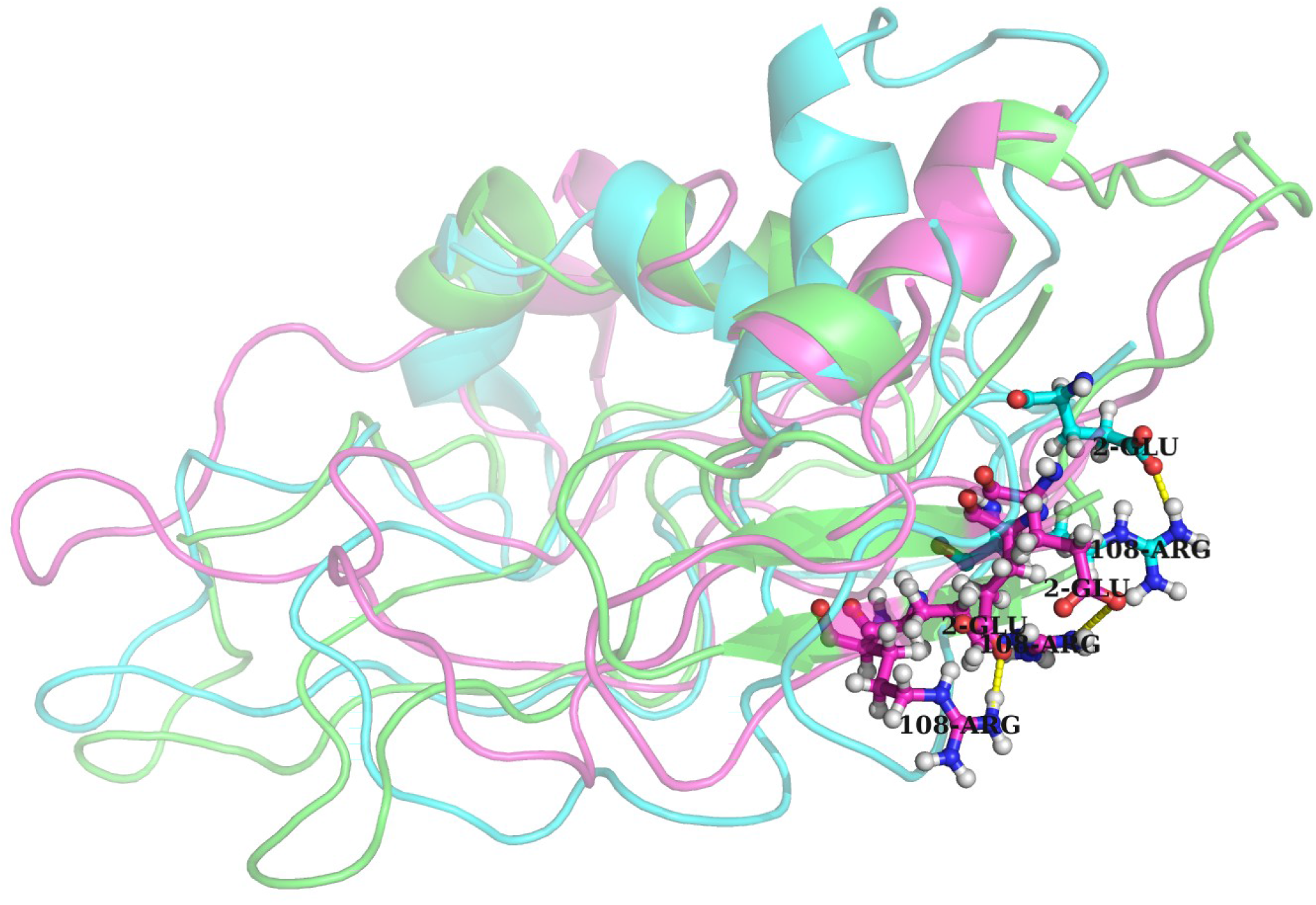
The long range persistent salt-bridge in 1CD3 demonstrating a case of a salt-bridge mediated long range secondary structural association.

The other long range persistent salt-bridge found in 1F0R (**Figure 7**) represented an exactly opposite case. Here the salt-bridge (41-Asp ~ 129-Lys) persisted only briefly *(pers:* 0.084: i.e., 8.4% of the time) with respect to the whole MD trajectory wherein it primarily supported two conformations (cluster-4, 5 with *pers:* 0.446, 0.133) persisting almost half the time in one of the two clusters (cluster-4) and one eighth in the other (cluster-5), while, its appearance in the rest of the clusters (cluster-1, 2, 3, 6) were virtually of flitting nature (*pers*: 0.002, 0.019, 0.018, 0.012). Clearly this is a representative case of conformational preference of a salt-bridge having a long range contact order (i.e., bringing together two far-apart regions of the disordered chain temporally for certain phases). Hence, this salt-bridge is exemplary to demonstrate the case of a representative salt-bridge particularly for cluster-4 in 1F0R.

**Figure 7.**
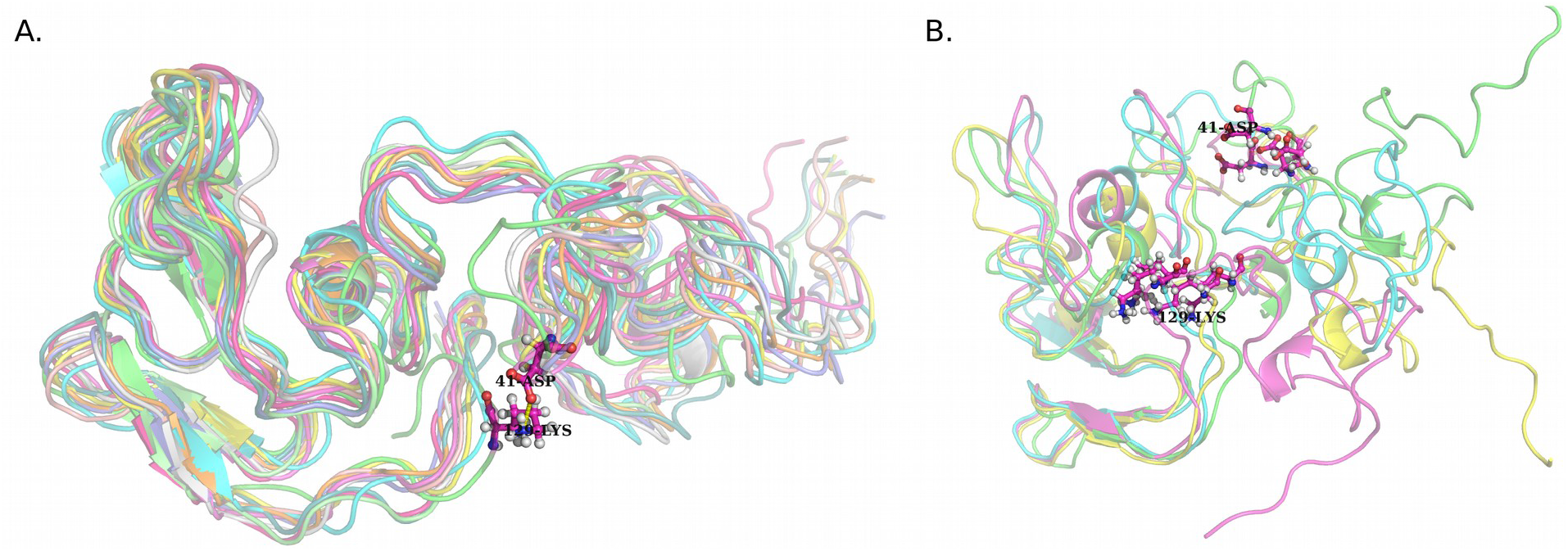
The long range persistent salt-bridge in 1F0R demonstrating a case of conformational preference of a salt-bridge having a long range contact order.

### 3.3. Representative Salt-bridges for conformational clusters

As standardized earlier [33], a persistence cut-off of 0.25 (25%) was used to define the ‘high’ persistent salt-bridges. Among the persistent salt-bridges found in each conformational cluster (for each protein) there were virtually two prototypes: (1) those persisted in all or most clusters (i.e., throughout the whole MD trajectory) and (2) those persisted in certain individual cluster(s). It naturally follows that the first prototype would give rise to high ‘overall’ persistence values (i.e., persistence calculated for the whole MD trajectory) which would generally decrease for the second. The first prototype of salt-bridges are therefore dynamically conserved in the idp, imparting general structural constraints, ‘common’ to all possible conformations while the second represents ‘unique’ temporal constraints particular to certain coformation(s).

As it turned out to be, the average persistence of salt-bridges computed cluster-wise (let’s call it cluster-persistence) were found to be much higher for the aforementioned ‘common’ prototype in comparison to the ‘unique’. A thorough statistics of the data further revealed that the average persistence of a salt-bridge generally increased with its cluster-occupancy (i.e., the number of confomational cluster the salt-bridge is found to be present in, with a ‘high’ persistence). This is perhaps reasonable, though not obvious, since, here, the analysis is based on cluster-persistence (i.e., persistence calculated per cluster), rather than overall persistence (i.e., persistence calculated for the whole trajectory). To elaborate the above point, let’s assume the case of a saltbridge found to be present throughout a particular cluster but absent otherwise across the (rest of the) MD trajectory. Such representative salt-bridges ‘unique’ to single conformational clusters would have retained really high cluster-persistence for the given cluster. In reality, the highest cluster-persistence for this category of salt-bridges (‘unique to a single cluster’) was found to be no more than 0.589 (for the salt-bridge 4-Lys ~ 9-Glu in 1F0R), followed by 0.560 (for 16-Lys ~ 22-Glu in Aβ42), and, 0.495 (for 32-Lys ~ 98-Asp in α-syn) while the average cluster-persistence was found to be 0.367 (±0.115) over 14 such ‘unique to single cluster’ instances of salt-bridges found across the four IDPs (**Table 1**). On the other hand, for salt-bridges found at high cluster-persistence among all clusters or throughout the whole molecular dynamic trajectory (the so called ‘common’ prototype) of salt-bridges, the same average was found to be 0.684 (±0.205), again, for 14 instances.

**Table 1.** Average persistence of representative salt-bridges for each conformational cluster (standard deviations given in parentheses) as a function of their cluster occupancy. Cluster Occupancy refers to the number of clusters a salt-bridge is found to be present in, with a high persistence (*pers* ≥ 0.25). The entries without a corresponding standard deviation refer to the ones with single occupancy (C1, C6 for Aβ42).

| Idp | Average cluster-Persistence as a function of Cluster Occupancy |  |  |  |  |  |
| --- | --- | --- | --- | --- | --- | --- |
| Cluster-number: | C1 | C2 | C3 | C4 | C5 | C6 |
| 1CD3 | 0.282<br>($\pm 0.040$) | 0.350<br>($\pm 0.063$) | 0.635<br>($\pm 0.225$) | - | - | - |
| 1F0R | 0.391<br>($\pm 0.118$) | 0.583<br>($\pm 0.124$) | 0.419<br>($\pm 0.090$) | 0.429<br>($\pm 0.117$) | 0.582<br>($\pm 0.166$) | 0.719<br>($\pm 0.173$) |
| α-syn | 0.403<br>($\pm 0.131$) | 0.329<br>($\pm 0.022$) | - | 0.430<br>($\pm 0.004$) | 0.704<br>($\pm 0.324$) | - |
| Aβ42 | 0.560 | 0.350<br>($\pm 0.119$) | - | - | - | 0.850 |

The degree of structural variation among the conformational clusters (or, in short, conformational variation) was estimated by the average of the pairwise root mean square deviation in C^α^ atoms between central conformations representative of each cluster. Overall, conformational variation among clusters in an idp was found to follow an inverse trend with the fraction of its ordered secondary structural content, also constrained by the number of representative salt-bridges in it. In other words, less number of salt-bridges and lesser degree of secondary structural content imparted more variation among the conformations. 1CD3, being the idp with the highest secondary structural content (56.7%) had the least variations (5.731 ±0.127 Å) among its three conformational clusters, constrained by 15 representative (7 common + 8 unique) salt-bridges. Furthermore for having two fairly long helices, the three clusters had clear visual resemblance in their overall shape wherein the variation indicated extensive movements of disordered loops connecting the helices, triggered and constrained by the representative saltbridges. Interestingly, in spite of being a partially disordered protein and having as many as 23 representative salt-bridges, 1F0R (among the four IDPs) exhibited the highest structural variation (10.850 ±3.00 Å) across its conformational clusters (also reflected visually, **Figure 8**). This apparently anomalous feature can be explained by the abundance of anti-parallel beta strands rather than helices as secondary structural elements in the protein chain resulting in a corresponding local clustering of the representative salt-bridges at different structural regions of the dynamic chain. It is a well known fact in protein science that proteins containing greater beta sheet content undergo far more severe deformations [48,49] than helical proteins, for beta sheets (and strands) are structurally less stable and geometrically less ordered than helices for more than one reason: (i) beta sheets get stabilized by inter-chain hydrogen bonding as compared to intrachain for helices and therefore are not self-sustainable like helices (ii) the influence of the backbone N-C^α^ -C (τ) bond-angle variation is much more pronounced in beta sheets compared to helices. For the completely disordered proteins, Aβ42 had a slightly higher degree of conformational variation (10.042 ±2.389 Å) than α-syn (8.500 ±1.840 Å). Aβ42 is constrained by a brief dynamic appearance of a small helix and only 4 representative salt-bridges, wherein, the conformations indicated open ended free movements of the overall protein chain, also contributed by its much smaller length (42 amino acids). On the other hand, α-syn did not give rise to any appreciable secondary structural presence in any of its conformations and were constrained by 9 representative salt-bridges. The much larger length (140 amino acids) of α-syn as compared to Aβ42 also potentially contributes to the lesser degree of variation in the former as it significantly enhances the influence of electrostatic interaction globally throughout the structure making the chain dynamically bent and concomitantly decreasing the scope and extent of open ended free movements (likewise to that of Aβ42).

**Figure 8.**
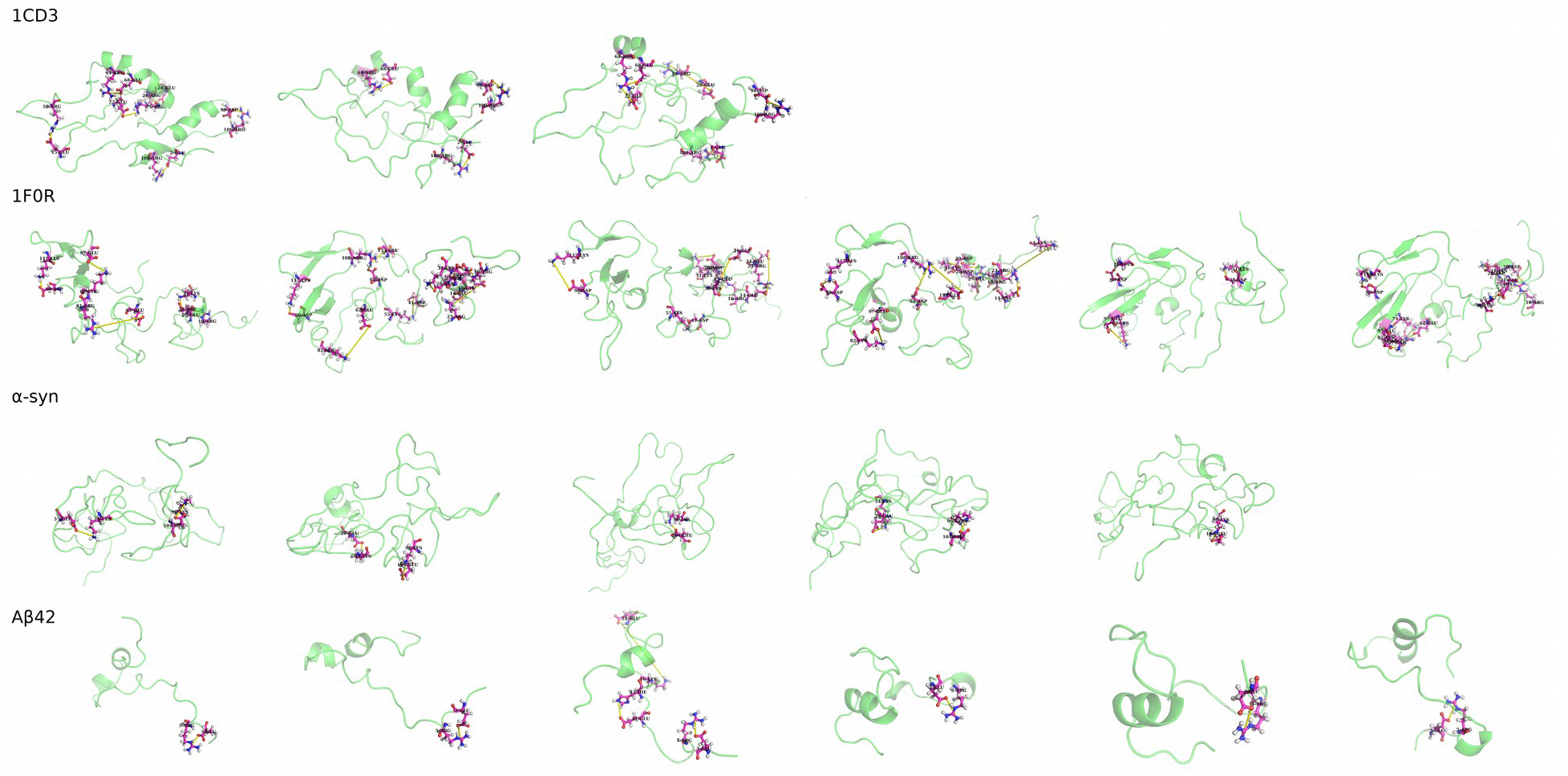
The conformational phases (presented by cluster centers) and their corresponding representative salt-bridges for the four IDPs (1CD3, 1F0R, α-syn, Aβ42 presented in rows 1-4 respectively).

### 3.4. Scale-freeness and Criticality in salt-bridge dynamics and phase transitions

As discussed vividly in the introduction, self-organized criticality (SOC) has often been characterized by scale-free distributions of appropriate representative parameters across physical, chemical, biological as well as other complex systems. In the current context of IDPs, it appears from the above analyses that salt-bridge formation (and the associated transient dynamics) can indeed be viewed as an indispensable aspect of the criticality associated to the complex phase transitions of these proteins among their structural conformations. In order to further verify the plausibility of the hypothesis, the distribution of ‘cluster persistence’ for the whole repertoire of salt-bridges (i.e., without using any persistence cut-off) were plotted in log-log plots. Interestingly, all the plots (without hitting a single exception) could best be fitted to straight-lines with descending (i.e., negative) slopes, and, thereby demonstrating power law distributions (y=k.x^-γ^) with lower order fractional exponents (**Figure 9, Figure S1-S4** in the **Supplementary Materials**). This also is a strong indication of the scale invariant self-similar fractal geometries in the structural conformations of these IDPs.

**Figure 9.**
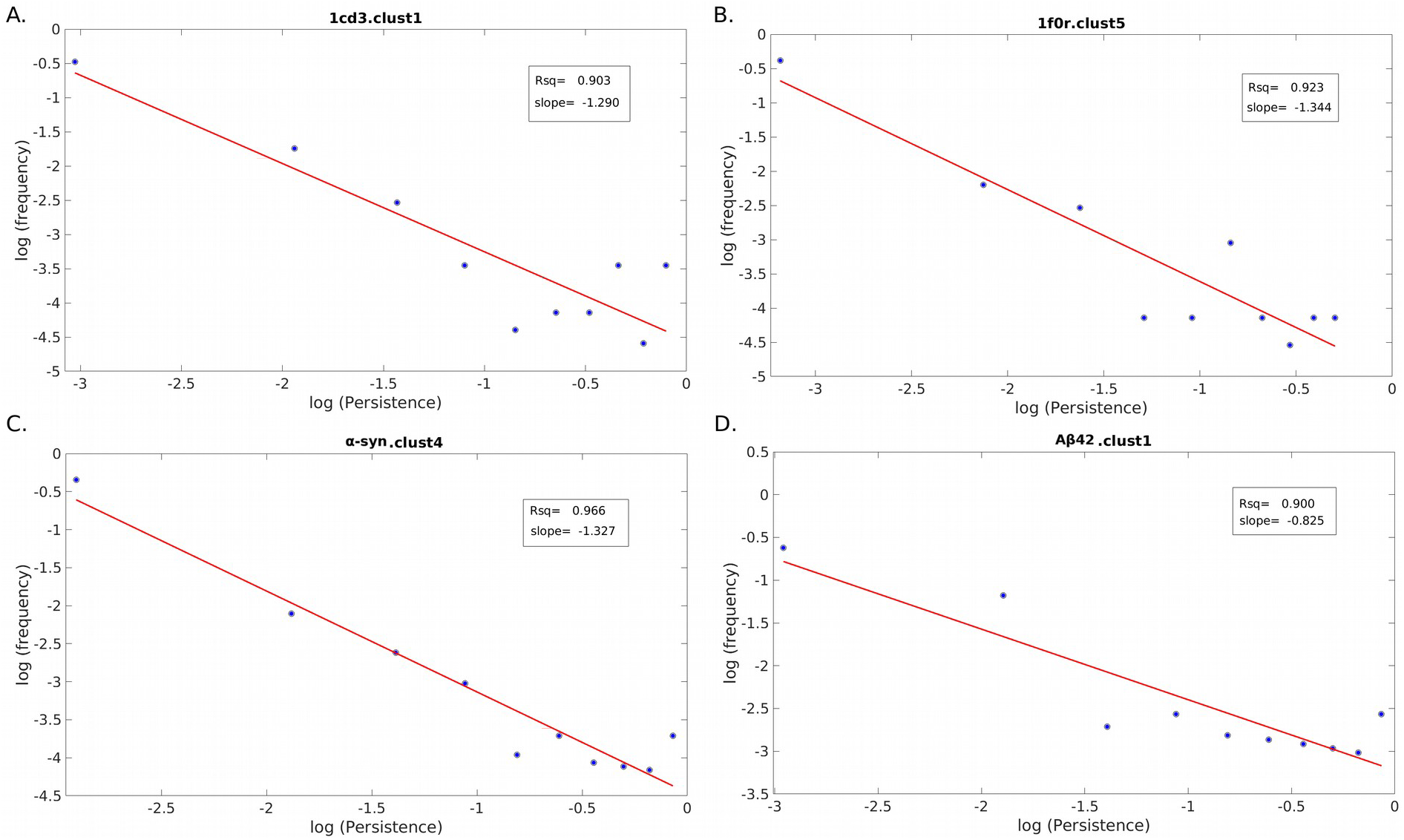
The log-log plot of the frequency versus persistence of salt bridges. Clusters with the highest absolute values of fractional exponents for the four proteins are plotted in A) 1CD3, B) 1F0R, C) α-syn & D) Aβ42.

In the log-log plots **(Figure S1-S4** in **Supplementary Materials)** of the frequency versus ‘cluster persistence’ of salt-bridges are plotted individually for the structural conformations (clusters) corresponding to 1CD3, 1F0R, α-syn, Aβ42 respectively followed by linear least-square fitting of the data. The least square fitted straight-lines are drawn in red. The associated goodness of fit were measured by the corresponding coefficients of determination (R^2^) which were found to be fairly high (averages over all clusters were 0.904, 0.800, 0.922, 0.811 respectively for 1CD3, 1F0R, α-syn, Aβ42) and statistically significant (p-values ≤ 0.05) as suggested by their corresponding p-values (average over the whole set: 0.0074 ±0.014). Hence, the frequency distributions of ‘cluster persistence’ of the salt-bridges were indeed found to carry signatures of power law distributions as is reflected from the values of the corresponding fractional exponents (i.e., the slopes of the corresponding best fitted straight-lines: −1.259, − 1.023, −1.26, −0.599 averaged over all clusters for 1CD3, 1F0R, α-syn, Aβ42 respectively).

The whole analysis gives us the essential insight that the transient nature of salt-bridge dynamics not only plays a pivotal role in retaining the overall desired confromational flexibility in IDPs (as was also revealed earlier [33]) but also further helps them to attain their quasi-stable conformational phases and traverse around these phases. The continuous transition among these conformational phases makes the IDPs behave like gels rather than crystalline solids to which the class of well folded (and perhaps more importantly well packed) globular proteins considerably resemble [50]. Therefore, in the current context of molecular dynamics of IDPs, the whole mechanism of salt-bridge formation can be envisaged equivalent to physical avalanche in sand-pile model [21] or neural avalanches [26,29], wherein, scale-freeness may potentially indicate emergence of self-organized criticality.

## 4. Conclusion

The primary objective of this paper was to analyze the structural disorder in IDPs and to find out the plausibility and extent of any hidden order based on self-similarity which might be responsible for their degenerate conformations. MD simulation of several IDPs reveal its different possible structural conformations among which the molecular structures get transformed in a self-organized fashion to allow the degeneracy and become intrinsically disordered. In this paper, different structural conformations of IDPs (as obtained in MD simulation) are cauterized in several groups based on their structural difference or RMS deviations. These structural groups or clusters are considered as dynamical phases in the molecular dynamics of corresponding IDPs. Hence, the phase transition dynamics are studied to analyze the persistence and long term behavior of the corresponding IDPs to be retained in the form of a bunch of representative structural conformations. It has also been revealed that the ‘transient dynamics’ of salt-bridges, which was earlier found to be a key to retain their structural flexibility [33], furthermore supports the aforementioned structural degeneracy and the proposed self similarity. Overall, the salt-bridges could be broadly classified into two groups: persistent and ephemeral. While persistent salt-bridges were found to be largely responsible in providing the desired structural formations to the corresponding IDPs even as they continuously undergo transitions among conformational clusters (phases), the ephemeral salt-bridges provided the essential conformational entropy among as well as within the clusters. The study also reveals that the transient dynamics of salt-bridges being a critical phenomenon in the molecular dynamics of IDPs allows these proteins to retain their criticality and complex phase transitions among different structural conformations. Also, it is observed that the overall distribution of persistence of salt-bridges in different structural phases could be characterized by power-law distributions with lower-order fractional exponents. This indicates scale-invariant self-similar fractal geometries in structural conformations of different IDPs. Thus, the salt-bridge dynamics could be compared with the avalanche mechanism in molecular dynamics. The scale-free behavior of saltbridge formation and dynamics might be an indication that IDPs are also retained around some critical points and allow themselves to transit between consecutive structural phases through selforganized criticality. The phase transition dynamics revealed that the structures, in the course of the equilibrium of the given IDPs resemble with one or two average structures (phases) for at least more than fifty percent of the ensembles. In itself, it is a strong insight in terms of understanding the overall structural degeneracy of the IDPs which may potentially facilitate structural tinkering, for example, in drug design for IDPs that are responsible for deadly neurodegenerative disorders. In more elaborate terms, each conformational cluster (or each structural phase) from a time-evolved IDP structure may be individually surveyed for its druggability or functional characterization computationally (e.g., protein-protein binding) – which would definitely aid benefits to the corresponding excercise.

## Acknowledgment and funding

The work was supported by the Department of Science and Technology – Science and Engineering Research Board (DST-SERB research grant PDF/2015/001079/LS).

## Footnotes

1 Systems that are intrinsically difficult to model due to their inherent dependencies and complexity of their interactions

2 For a period of time / time dependent / time-related – used contextually

3 Ideal chains in polymer science are characterized by a theoretical shape factor of 1.5

